# Linearizing and forecasting: a reservoir computing route to digital twins of the brain

**DOI:** 10.1101/2024.10.22.619672

**Authors:** Gabriele Di Antonio, Tommaso Gili, Andrea Gabrielli, Maurizio Mattia

**Affiliations:** “Enrico Fermi” Research Center - CREF, 00184 Rome, Italy; Dip. di Ingegneria Civile, Informatica e delle Tecnologie Aeronautiche, Università degli Studi “Roma Tre”, 00146 Rome, Italy; Natl. Center for Radiation Protection and Computational Physics, Istituto Superiore di Sanità, 00161 Rome, Italy; Networks Unit, IMT Scuola Alti Studi Lucca, 55100 Lucca, Italy

## Abstract

Exploring the dynamics of a complex system, such as the human brain, poses significant challenges due to inherent uncertainties and limited data. In this study, we enhance the capabilities of noisy linear recurrent neural networks (lRNNs) within the reservoir computing framework, demonstrating their effectiveness in creating autonomous *in silico* replicas – digital-twins – of brain activity. Our findings reveal that the poles of the Laplace transform of high-dimensional inferred lRNNs are directly linked to the spectral properties of observed systems and to the kernels of auto-regressive models. Applying this theoretical framework to resting-state fMRI, we successfully predict and decompose BOLD signals into spatiotemporal modes of a low-dimensional latent state space confined around a single equilibrium point. lRNNs provide an interpretable proxy for clustering among subjects and different brain areas. This adaptable digital-twin framework not only enables virtual experiments but also offers computational efficiency for real-time learning, highlighting its potential for personalized medicine and intervention strategies.

## Introduction

The study of complex dynamical systems has a central role in contemporary scientific research. Over recent decades, a plethora of data-driven methodologies has emerged, denoting a field in rapid evolution that is capable to addressing systems with increasing complexity. Among these methodologies, machine learning techniques have become increasingly popular due to their effectiveness in modeling complex datasets and deliver accurate predictions (1, 2, 3).

Within the spectrum of machine learning techniques, reservoir computing (RC) has distinguished itself as a powerful approach for processing temporal data. RC leverages high-dimensional recurrent neural networks (RNNs) with fixed internal couplings to transform input time series into rich, high-dimensional representations (4, 5). This transformation enables the prediction of future samples by simply reading out the current state of the network, effectively capturing the system’s dynamics (6, 7, 8). The elegance of this approach lies in conceptualizing the readout as a projection onto a manifold learned through linear regression, echoing the efficient coding strategies observed in biological neural networks (9, 10, 11). Consequently, RNNs have become invaluable tools in neuroscience, aiding in hypothesis generation and providing analytical frameworks for understanding neural computations (12, 13, 14, 15).

However, as machine learning has become increasingly popular, some clarity of understanding may have been lost in the pursuit of better predictions. While the inferred models are often quite accurate, they can sometimes lack interpretability (16, 17, 18). A valuable framework to address this limitation is provided by Koopman theory, which offers clear insights into the system dynamics by mapping nonlinear systems into simple yet higher-dimensional dynamics (19, 20, 21, 22). This approach involves the development of data-driven methods aimed at computing finite-dimensional linear representations of nonlinear systems within a suited functional space. Once an equivalent linear system is inferred, interpretability becomes straightforward, allowing to gain understanding of the time scales and dynamical modes of the observed systems.

The capability to predict future states of a dynamical system based on past observations allows, in principle, to build a replica of the same system. Indeed, predicted observations can be fed back as input resulting in a generative model. Both Koopman-based methods (Fig. 1A) and RC approaches (Fig. 1B) have the potential to generate these digital copies by producing time series statistically equivalent to those generated by the original systems. The inferred generative models can then serve as ‘digital twins’ – a concept that has emerged as a transformative paradigm in the study of dynamical and natural systems (23, 24). Such digital twins can be effective tools for analyzing and simulating their physical counterparts. Successful examples include digital copies of brain activity designed to assist neurosurgeons in dealing with drug-resistant epileptic patients (25, 26, 27). Furthermore, these tools can provide a complementary workbench for designing stimulation approaches to be tested *in silico*, ultimately leading to control systems for neurorehabilitation (28).

**Figure 1.**
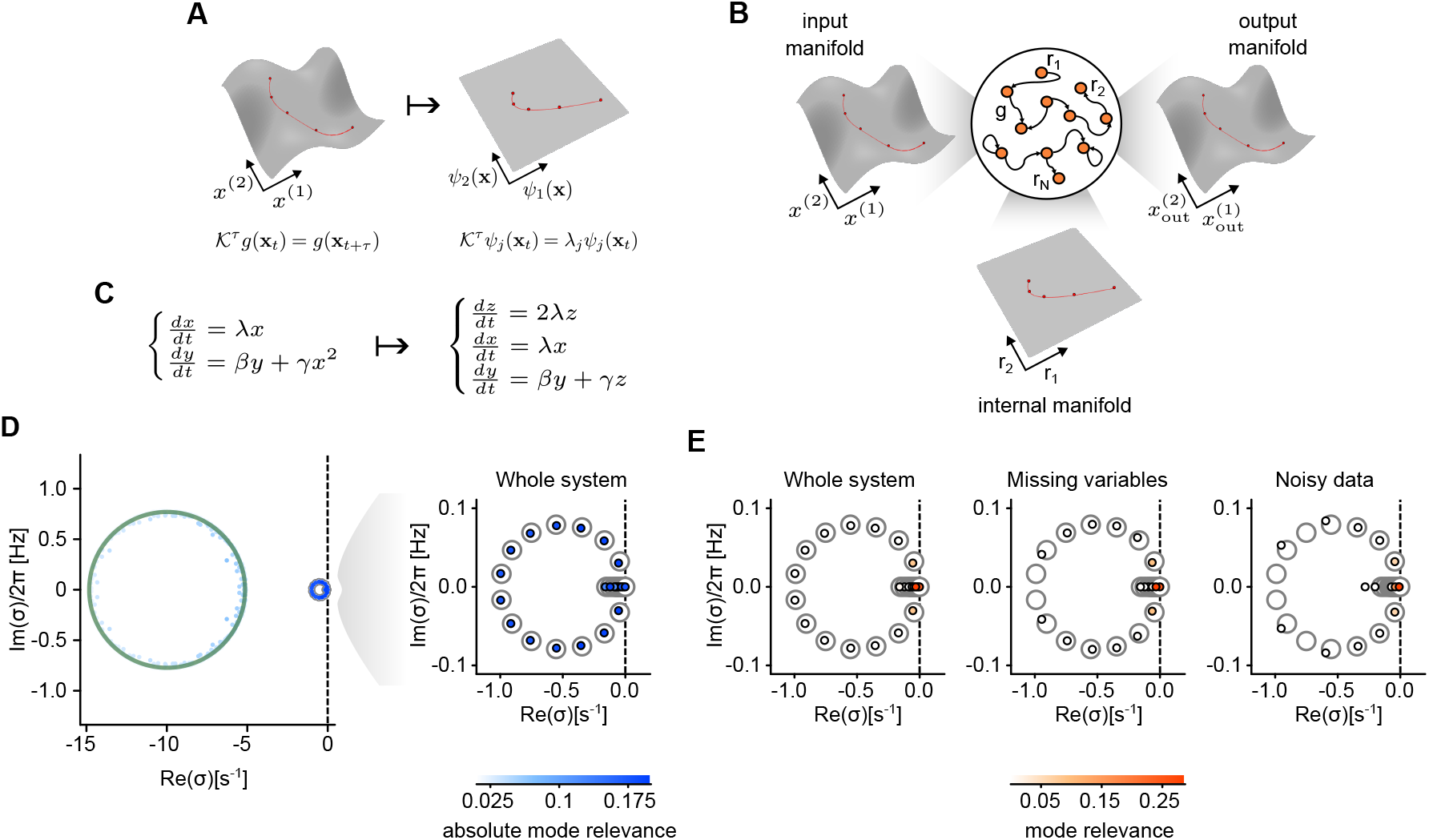
lRNN encodes a linear representation of the input system. (**A**) Nonlinear dynamics can be projected onto linear manifolds. Within the infinite functional space of the system observables, the eigenfunctions of the Koopman operator establish a basis for decomposing nonlinear dynamics into simpler linear processes. (**B**) Reservoir computing (RC) with recurrent neural networks (RNNs). After ‘learning’, the readout weights ***W*** ^out^ are set to reproduce the system observables provided as input. The resulting network is composed of linear units (lRNN) and its post-learning autonomous dynamics encapsulates a linear representation of the original system. (**C**) Example of a 2-dimensional nonlinear system that can be decomposed into a set of 3 linearly evolving variables. (**D**) Spectrum of a lRNN with *N* = 500 units trained to mimic a 15-dimensional version of the system in (C) composed of 5 linear *x* variables and 10 nonlinear *y* variables. Green circles, spectrum of the lRNN before training. Gray circles, theoretically-derived eigenvalues of the 20-dimensional linearized system to be replicated (see text for details). Small circles, Laplace transform poles from the post-learning autonomous lRNN replicating the example system. Filling colors code the absolute mode relevance ⟨ |Ξ|_*jk*_|⟩_*j*_. Right panel, zoom in of the left panel showing the 20 most significant modes. (**E**) Same spectrum as in (D) with filling colors coding the mode relevance ⟨|Ξ_*jk*_*v*_*k*_(0)|⟩_*j*_ (left panel, see text for details). Central panel, spectrum of a lRNN trained on incomplete information (i.e., omitting all *x* and 5 *y* variables). Right panel, spectrum of a lRNN trained on the same observables but perturbed by noise.

In this study, we investigate the potential of RNNs with linear units to implement effective digital twins of single-subject brain dynamics, measured as the vascular response to neuronal activity (i.e., the BOLD signal) extracted from functional magnetic resonance imaging (fMRI). We demonstrate the autoregressive nature of this approach and its effectiveness in representing whole-brain activity in a low-dimensional latent state space. Such a reduced representational complexity mitigates the risk of overfitting often associated with one-to-one fitted physical models, especially in realistic scenarios where only a limited number of observations are available and only a partial view of the system can be experimentally accessed. The integration of endogenous memoryless fluctuations is another essential component that enables the digital twin to self-sustain a noisy dynamics statistically equivalent to the one experimentally measured.

## Results

### Linear RNNs can copy nonlinear systems

We start investigating the reservoir computing abilities of a RNN with *N* linear units (lRNN). The state of the lRNN is the vector ***r***(*t*) whose elements are the activities *r*_*i*_(*t*) of each unit, following the linear dynamics

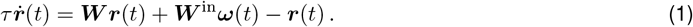

The synaptic matrix ***W*** ∈ ℝ^*N ×N*^ determines the recurrent input ***W r***, which, together with the contributions from the external environment ***W*** ^in^***ω***(*t*), composes the total input ***h*** that each unit receives. In general, these currents are nonlinearly transformed by an input-output gain function Φ(***h***) we assume here to be Φ(***h***) = ***h*** – a condition usually associated with unit activities perturbatively fluctuating around the quiescent state (see Materials and Methods). Network units receive a source of external input due to the *M* observables *ω*_*i*_(*t*) of the systems under investigation, mediated by the synaptic matrix ***W*** ^in^ ∈ ℝ^*N ×M*^. From Eq. (1) the network state at any time *t* results then to be

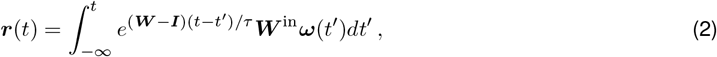

where ***I*** ∈ ℝ^*N ×N*^ is the identity matrix.

If the lRNN is capable to display the so-called ‘generalized synchronization’ property in response to the input ***W*** ^in^***ω*** (29, 30), ***r***(*t*) effectively embeds the dynamics of the ‘whole’ observed system into its state space. This because the lRNN incorporates the ‘echo’ (31, 4, 32) of past observations implementing a dimensional embedding *a la* Takens (33, 34, 35). In this RC framework, for sufficiently high *N* a simple linear transformation capable to approximately map the network state to a target output exists

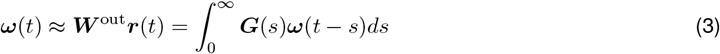

with synaptic weights ***W*** ^out^ ∈ ℝ^*M ×N*^ computed via a simple linear regression (4, 5). The linear kernels ***G***(*s*) = ***W*** ^out^*E*^(***W*** *−****I***)(*s*)*/τ*^ ***W*** ^in^ ∈ ℝ^*M ×M*^ (i.e., the Green functions) resulting from Eq. (2) reveals the autoregressive nature of the RC approach in this lRNN case. This concept can be better clarified by resorting to the eigendecomposition of the synaptic matrix ***W*** = ***Q*Λ*Q***^*−*1^. Here **Λ** is the diagonal matrix of the eigenvalues

(Λ_*ii*_ = *λ*_*i*_) and the eigenvectors are the columns of ***Q***: ***W***|*Q*_*i*_⟩ = *λ*_*i*_|*Q*_*i*_⟩. By applying this decomposition to the kernel expression, a superposition of *N* exponential modes results:

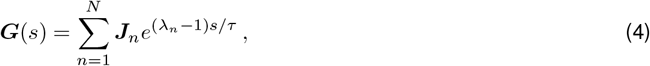

where 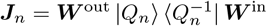 are *N* rank-1 matrices with 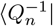 being the *n*-th row of the inverse matrix ***Q***^*−*1^ (see Materials and Methods). The convolution in Eq. (3) is integrable if the spectral radius *ρ* = max_*n∈*[1,*N*]_ Re *λ*_*n*_ of ***W*** is smaller than 1. Under this condition, unperturbed lRNNs (***ω*** = **0**) have a stable equilibrium point at ***r*** = **0**, and self-consistency equation (3) describes a continuous-time autoregressive (AR) model with order *N* (number of exponential modes in ***G***(*s*)). This relationship between RC with lRNNs and AR models generalizes previous results obtained in the discrete-time domain (36, 37). As we will see later, the order of the equivalent AR model can be much smaller than *N* as many ***J***_*n*_ are usually close to **0**.

By replacing the observables ***ω***(*t*) provided as input with the reconstructed once from Eq. (3), the lRNN results to follow the autonomous dynamics

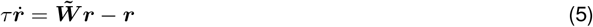

where 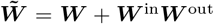 is the updated (i.e., ‘learned’) synaptic matrix. If the network can correctly predict the future of the target output, Eq. (5) provides a linear representation of the system under analysis. Relying as above on the eigenmode decomposition 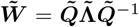, the projections 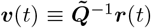 evolve in time as a set of decoupled variables:

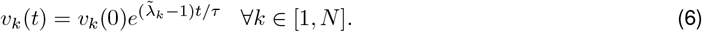

This last step makes it apparent the relationship with the Koopman-operator theory, as a suited linear combination of these modes eventually implements an effective decomposition of the time series ***ω***(*t*):

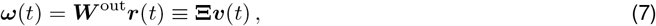

with 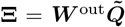. The Laplace transform of the reconstructed observable will then have *N* poles 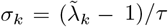 with their related residues, allowing us to represent the dynamical properties of the ‘digital twin’ (i.e., the autonomous lRNN), and linking it to AR models and finite approximations of the Koopman operator (see Materials and Methods). Indeed, the *k*-th residue is proportional to Ξ_*jk*_*v*_*k*_(0) and provides in absolute value the relevance of the *k*-th mode in describing the observable *ω*_*j*_, given the initial network state ***r***(0).

As a testing ground for the theoretical framework derived above we consider the following nonlinear system:

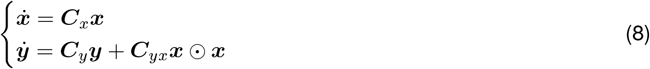

Choosing ***C***_*x*_ as a diagonal matrix, this system can be mapped onto a linear one by introducing a set of new observables ***z*** = ***x*** ⊙ ***x*** = **diag**(***x***) ***x*** (Fig. 1C). In this case, the related infinitesimal generator of the Koopman operators is equivalent to the finite matrix ***K*** with diagonal blocks ***C***_*x*_, 2***C***_*x*_, ***C***_*y*_.

We trained a lRNN to replicate the relaxation dynamics of this system from the observables ***ω*** = [***x***; ***y***] with ***x ∈*** ℝ^5^ and ***y ∈*** ℝ^10^, starting from a random initial condition ***ω***(0). As initial synaptic matrix ***W*** we chosen the one associated to ring-like connectivity (i.e., *W*_*i,i*+1_ = *W*_*N*,1_ = *ρ* are the only non-zero elements) known to have optimal performances in the case of linear units (38). In Fig. 1D the poles *σ*_*k*_ of the reconstructed observables from Eq. (7) are shown together with the related mode relevance (color intensity). Despite the nonlinearity of the system to replicate, the overlap between the poles *σ*_*k*_ and the theoretical spectrum of ***K*** is remarkable. Furthermore, only 20 out of the *N* = 500 modes have ⟨|Ξ_*jk*_|⟩_*j*_ significantly different from 0 (blue-colored circles).

Decomposition modes display damped oscillations as 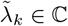 and a few of them, the slowest ones, contribute significantly to the dynamics (Fig. 1E, red-colored small circles). Such low-dimensionality is not altered even if during learning the lRNN receives noisy-contaminated observables or some of them are not provided as input. Indeed, the autonomous lRNN continues to predict accurately the system’s future also under these conditions, although the less relevant poles move away from the imaginary axis (missing match between small and large circles).

### lRNNs successfully forecast fMRI time series

To explore the applicability of the RC with lRNN, we examined time series data from a real-world physical system. We considered the brain activity of 20 healthy subjects at rest (i.e., in a state of quiet wakefulness), measured through fMRI. The experimental time series was derived from the BOLD signals recorded from a subset of voxels, which serve as proxies for neuronal activity across 11 cortical areas in the language network (Fig. 2A, see Materials and Methods).

**Figure 2.**
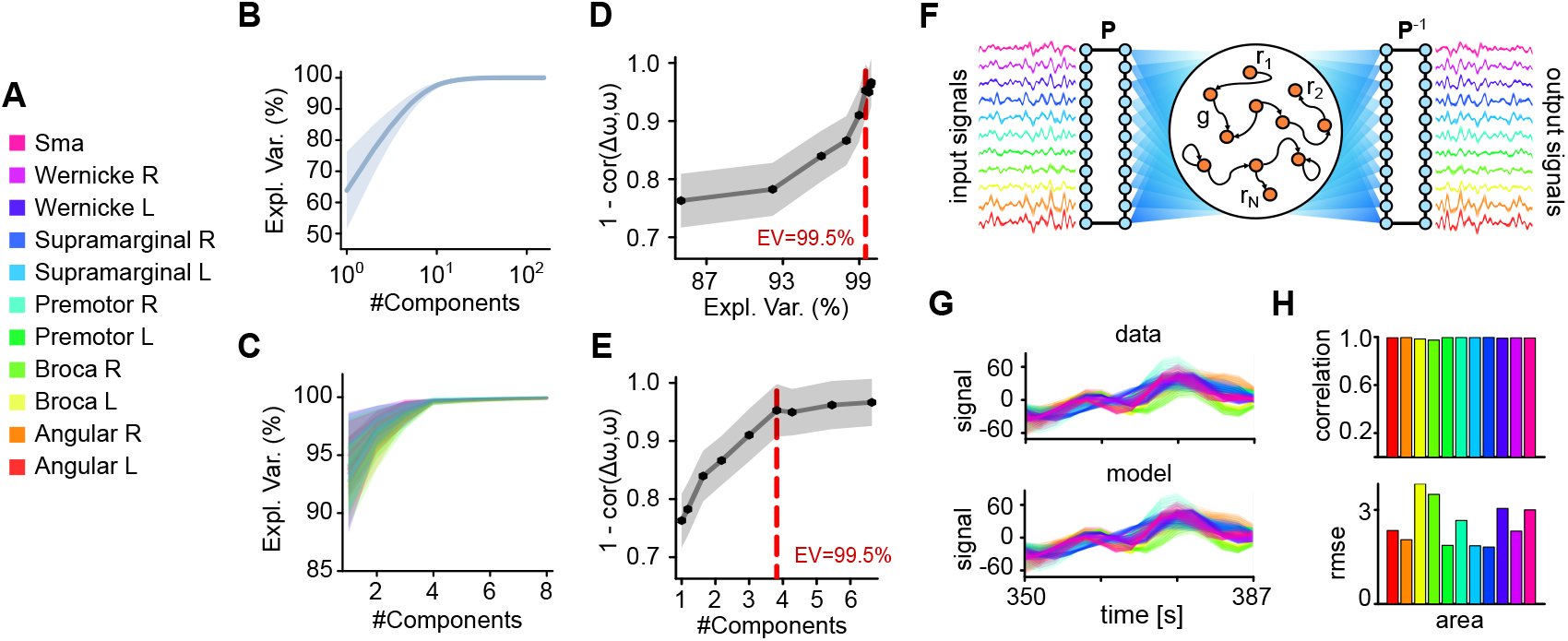
Compression and prediction of resting state fMRI activity. (**A**) Color legend associated with the 11 cortical areas taken from the language network. (**B**) Explained variance (EV) from PCA of the fMRI BOLD signals of the voxels in the language network. Solid line, average across the 20 considered subjects. Shaded area, standard deviation across subjects. (**C**) EV of the PCA for each cortical area. Colors as in (A). (**D** and **E**) 1 minus the correlation between the residuals from the PCA reconstruction (Δ*ω*) and the original data (*ω*) per single area in the last (unseen) 37.5 seconds. A value equal to 1 means that the remaining information is just white noise. Vertical dashed lines, explained variance (D) and number of first PCs (E) required to reach the EV of 99.5%. (**F**) Schematic design of the used lRNN. The input from each area is compressed to have an EV of 99.5%. ***P*** is the matrix operating this linear transform leading to an average of 44 observables. The lRNN output reconstructing these observable is then uncompressed applying ***P*** ^*−*1^. (**G**) Example of forecasted (bottom) and original (top) activities during the test window (last 37.5 seconds). Each curve is the BOLD signal of a voxel with color indicating the cortical area of belonging. (**H**) Forecast performances of the post-learning autonomous lRNN per area. Correlations between predicted and experimental activity (top) and root mean square error (bottom).

The BOLD time series underwent preprocessing using principal component analysis (PCA) to reduce redundancy and achieve dimensionality reduction, which was found to be relatively high (see Fig. 2B, C). Specifically, we assessed the correlation between the information lost during dimensionality reduction and the actual signals. Our analysis revealed that the first principal components (PCs), which explain 99.5% of the variance (Fig. 2D,E), excluded a residual activity that was indistinguishable from white noise. Based on this criterion, we determined that, on average, only four PCs per cortical area should be considered. Consequently, the average dimensionality of the dataset was *M* ≈ 44 PCs per subject (refer to Materials and Methods for details). This relatively high-dimensional time series served as the input for the lRNN. The readout synaptic matrix ***W*** ^out^ was computed over the first 280 seconds (Fig. 2F).

The decision to utilize a transformation of brain activity that results in information loss may appear as counter- productive. However, projecting onto the first PCs was expected to be less sensitivity to the uncertainties inherent in the measured BOLD signals. By employing these first PCs as inputs, we provided the lRNN with denoised representations of brain activity. Moreover, thanks to the echo state property (31, 4, 32), the missing information regarding the state of the observed system can be effectively recovered through the dimensional embedding that the lRNN performs.

As previously described, with the learned ***W*** ^out^ – which varies from subject to subject – we established a closed loop between the input and output of the lRNN, effectively creating an autonomous digital twin of the brain network under investigation. This approach yielded a remarkable overlap between the reconstructed and measured time series (see Fig. 2G for an example subject). At the population level, we evaluated the forecasting performance by calculating the correlations between replicated and experimental BOLD signals, as well as the root mean square errors (RMSEs) for the final unseen 37.5 seconds of the time series (Fig. 2H). From this perspective, the inferred digital twin demonstrated high-quality reconstruction of brain activity, further validating its effectiveness.

### lRNN as a digital twin of brain activity

The lRNN dynamics described thus far is dissipative, meaning that the state of the network ultimately relaxes to **0**. However, brain activity associated to the BOLD signals fluctuates without rest. To reconcile this discrepancy in our digital twin, we incorporated an endogenous noise into each network unit. This modification introduces a generative mechanism that sustains activity over time. More specifically, the autonomous dynamics of the inferred (i.e., learned) lRNN dynamics is a multi-variate Ornstein-Uhlenbeck process

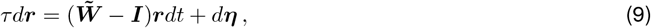

where ***η***(*t*) is the the vector of Gaussian white noise each unit of the network independently receives (⟨*dη*_*j*_(*t*)*dη*_*k*_(*t*^*′*^⟩) = *γ*^2^*δ*_*jk*_*δ*(*t* − *t*^*′*^)) with noise intensity *γ*. Note that, since the system is linear, this formulation does not change the theoretical framework derived above; in this case, it applies to the expectation values 𝔼[***r***] and 𝔼[***ω***] = ***W*** ^out^𝔼[***r***].

To validate the effectiveness of this stochastic version of the lRNN in accurately replicating brain activity, we numerically integrated the network dynamics for an additional 387.5 seconds (Fig. 3A). We then compared the functional connectivity (FC) – defined as the correlation between BOLD signals of all possible voxel pairs – calculated from this simulation to that obtained from the experimental data (Fig. 3B). Furthermore, the temporal evolution of the FC, referred to as functional connectivity dynamics (FCD), closely aligned with the experimental observations. This similarity was quantified by the Jensen-Shannon distance (JSd) between the distribution values of the associated matrices (Fig. 3C).

**Figure 3.**
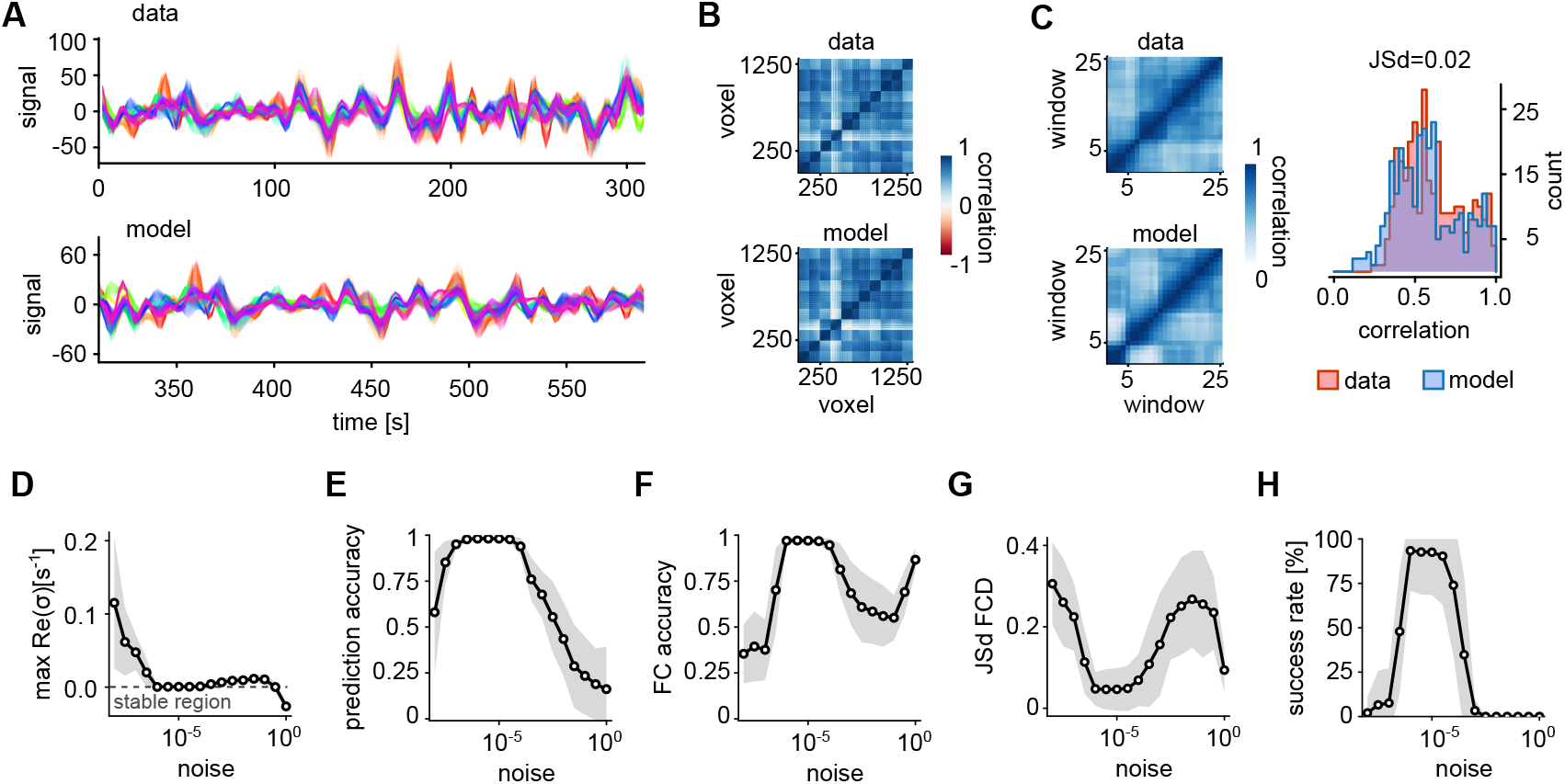
Endogenous noise in lRNN is needed to replicate brain activity. (**A**) Comparison of actual brain activity (top) with the corresponding output from a trained autonomous lRNN over the subsequent prediction interval (bottom). (**B** and **C**) Analysis of functional connectivity (FC) and functional connectivity dynamics (FCD) computed from the activity depicted in (A). Right, distribution of FCD values for lRNN-generated and exeprimental time series with the related Jensen-Shannon distance. (**D, E, F** and **G**) Features and performance statistics of inferred lRNNs across subjects varying the amplitude of endogenous noise: maximum pole real part (D), correlation between data and model prediction (E), FC correlation with actual data (F), and Jensen-Shannon distance between FCDs (G). (**H**) Success rate in reproducing above-threshold performances of the features in panels (E-G) (see main text for details).

Finally, we conducted a comprehensive analysis of the optimal intensity of noise across all subjects (Fig. 3), focusing on the accuracy of BOLD signal forecasting and the lRNN capability to reproduce both FC and FCD. To evaluate the quality of these aspects simultaneously, we introduced a success rate defined by applying fixed thresholds to the performance metrics: accuracy *>* 80%, FC correlation *>* 85% and JSd *<* 0.1. Additionally, we assessed the stability of the inferred lRNN by examining the maximum real part of the poles *σ*_*k*_ (Fig. 3D). This analysis enabled us to identify a common optimal intensity *γ* of endogenous noise across subjects with a standard deviation of 10^*−*6^ (Fig. 3h). Interestingly, the existence of an optimal noise displays some similarities with the resonance phenomenon found in other machine-learning studies (39), which in our case robustly emerged in all the subjects.

### Dynamical properties of the digital twin

We showed that RC with lRNNs can effectively reproduce resting state fMRI activity. Each simulated voxel can be described as a decomposition of linearly evolving modes. The dynamical properties of this decomposition can be effectively represented by the poles *σ*_*k*_ of the Laplace transform of the inferred lRNN, which are the eigenvalues 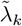 of the learned synaptic matrix 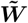. This spectrum of eigenvalues is shown in Fig. 4A for an example subject. For each of the modes, the relevance is averaged across all voxels and it is color coded, showing a pattern similar to what seen in Fig. 1E where only a subset of poles are moved towards the imaginary axis. These poles are the most relevant and display a specific organization like a rotated parabola. The modes with the highest relevance (i.e., the darkest) appear to be distributed at specific frequencies falling into the range of infra-slow oscillations (*<* 0.1 Hz), a typical footprint of the resting state and of the unconscious brain activity (40, 41, 42, 43). As a representative example, in Fig. 4A, the most relevant mode is one of the most persistent, characterized by a relatively small Re *σ*_*k*_, and exhibits a resonant frequency of approximately 0.02 Hz.

**Figure 4.**
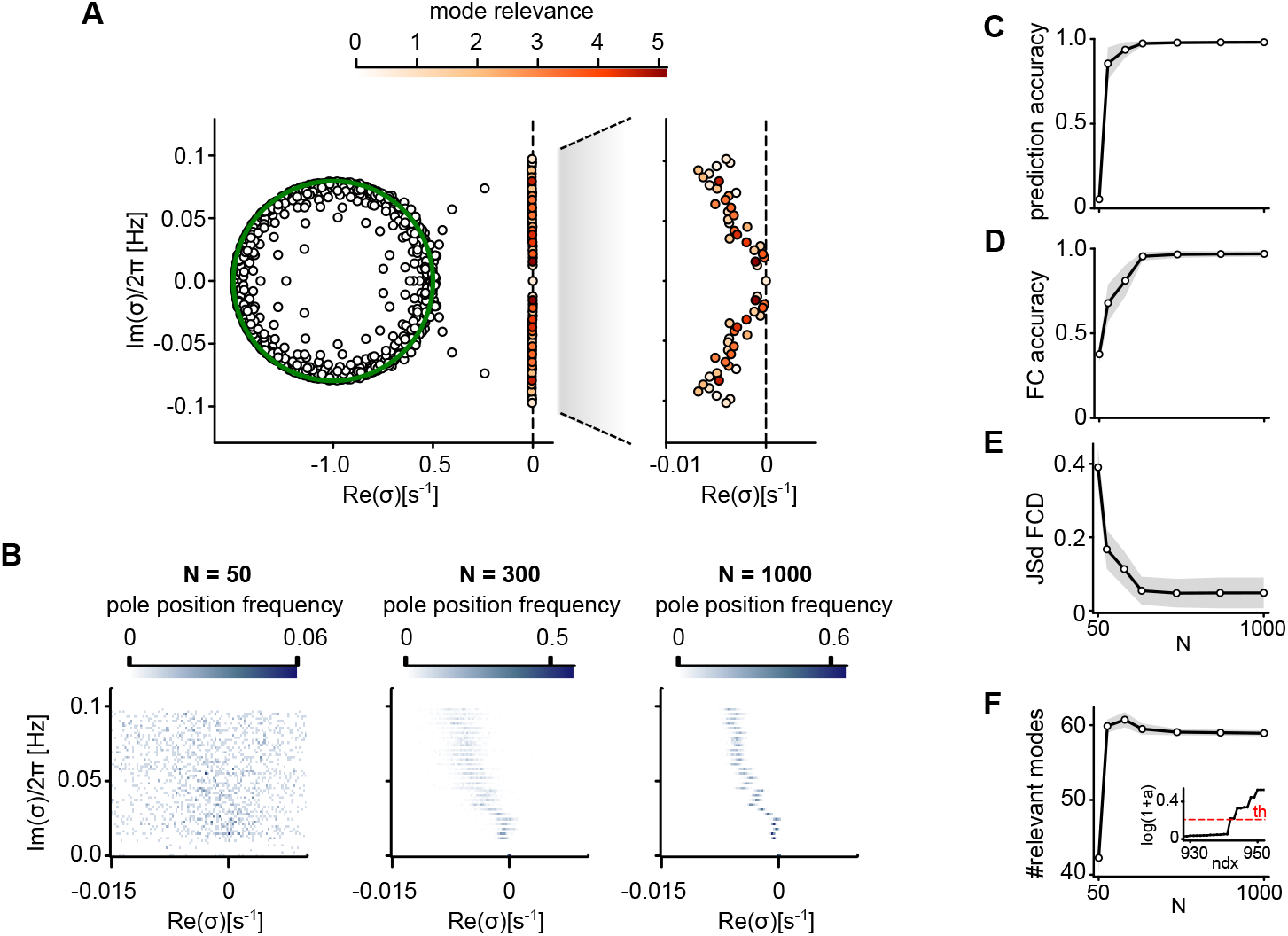
Convergence of the mode decomposition with increasing lRNN size. (**A**) Mode decomposition of a single cortical area for an example subject. Eigenvalues of 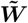 are color-coded as in Fig. 1E. (**B**) Distribution of eigenvalues for three distinct network sizes *N*, analyzed across 100 randomly sampled lRNNs. (**C**) Number of relevant modes (i.e., with relevance greater than 5%). Inset, sorted mode relevance and related threshold (dashed line). (**D**) Accuracy of forecasted activity. (**E** and **F**) Reproduction fidelity of FC (E) and of FCD (F). Gray shadings and black lines, standard deviation and mean across subjects, respectively.

Subsequently, we investigated the stability of this representation exploring how the spectrum of 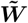 and the mode relevance change according to the number *N* of units in the lRNN. As shown in Fig. 4B, the density of eigenvalues in the complex plane (Re *σ*, Im *σ*) becomes less and less scattered with increasing network size. Besides, all the performance measures converged to fixed values corresponding to a high quality of the data replicate by the digital twin (Fig. 4C-E), proving that a finite size lRNN can reach asymptotically high performances. Intriguingly, as convergence is approached the average number of relevant modes stands at about 60 independent components (Fig. 4F). This number is significantly higher than the number of experimental PCs provided as input to the lRNN (44 on average). Thus, brain activity replicated by the inferred lRNN appear to live in a latent state space whose dimensionality is larger than the one determined by its observation.

Functional connectivity illustrated in Fig. 5A-top for an example subject is then fully replicated by the inferred (and stochastic) lRNN. This effective copy occurs even though the dimensionality of the latent state space of the digital twin is significantly smaller than the number of voxels encompassing the examined language network. To gain a deeper understanding of this result, we ‘opened the box’ by examining the linear transformation ***Z*** ≡ ***P*** ^*−*1^**Ξ ∈** ℝ^*L×N*^. According to Eq. (7), this matrix facilitates the mapping of the *N* eigenmode projections ***v***(*t*) onto the *L*-dimensional voxel-wise BOLD activity, represented as ***x***(*t*) = ***Zv***(*t*). The absolute values of the elements in this matrix are displayed in Fig. 5B for the same subject. Unsurprisingly, the most significant contributions arise from the slowest (rightmost) modes, specifically those with the largest Re 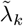. Additionally, by measuring the covariance between the eigenmode projections *v*_*k*_(*t*), it becomes evident from Fig. 5C that they are largely uncorrelated. Only about 60 of the slowest modes exhibit significant variability, as indicated by the dark diagonal elements. This empirical evidence suggests that functional connectivity can be estimated directly using the cosine similarity S_C_ between the rows of the matrix ***Z*** (see Materials and Methods):

**Figure 5.**
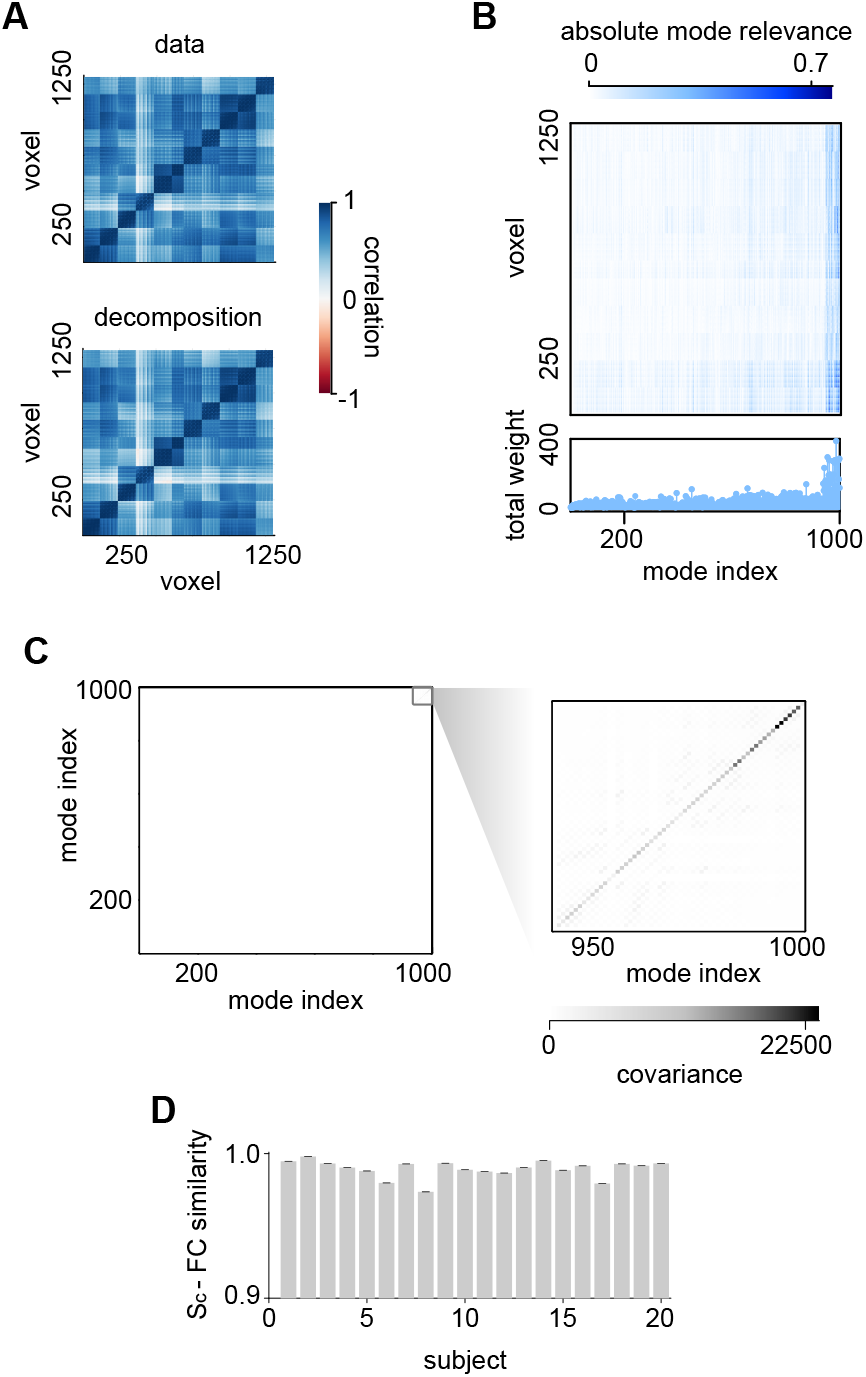
Low-dimensional mode decomposition explains functional connectivity. (**A**) Functional connectivity of an example subject (top) and cosine similarity (bottom) between rows of the matrix ***Z*** mapping eigenmode projections to voxel-wise BOLD activity from the inferred lRNN with *N* = 1000 units (see main text). (**B**) Absolute mode relevance |*Z*_*jk*_| for the same lRNN together with the total weight Σ_*j*_ |*Z*_*jk*_| of each of the *N* modes (bottom). Modes are sorted according to the real part of the associated eigenvalue (Re 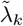). (**C**) Covariance matrix of the eigenmode projections *v*_*k*_(*t*) (see Eq. (7)) during the extended autonomous phase (387.5 seconds post learning in Fig. 3A-bottom). Right, zoomed-in view on the slow and most relevant modes. (**D**) Correlation between the upper diagonal elements of the FC and the cosine similarity (S_C_) of the rows of ***Z*** for each subject. Averaged over 100 noise realizations. Error bars, SEM.

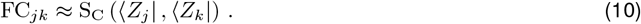

In Fig. 5A-bottom, the similarity matrix is presented, showcasing a remarkable overlap with the experimental functional connectivity **FC**. It is important to note that in computing ***Z***, we considered only the columns of **Ξ** associated with the relevant modes. This further confirms that the remaining *N ™* 60 degrees of freedom are nearly irrelevant.

### lRNN as a proxy to characterize subjects and brain areas

Given that inferred lRNNs reliably replicate observed brain activity and their modal decomposition defines their dynamical properties, a critical question emerges: Can lRNNs serve as proxies for understanding the similarities and differences among subjects and cortical areas? To address this, we characterized the spectrum of relevant eigenvalues by focusing on two key features: the linear relationship between their real and imaginary parts, and the overall significance of each oscillation frequency, as illustrated in Fig. 6A. The slope of this linear relationship indicates the correlation between decay times and oscillation frequencies, while the coefficient of determination (*R*^2^) assesses the goodness of fit for this linear model. In this two-dimensional parameter space, subjects are systematically distributed among three distinct regions (Fig. 6B). Those with high *R*^2^ values display either steep or shallow slopes, whereas subjects with low *R*^2^ values indicate a poor linear fit and suggest the presence of persistent isolated high-frequency oscillatory modes. Within this representational space, spectra characterized by steeper slopes (located at the bottom) are associated with poles that are closer to the imaginary axis. This suggests that those lRNNs (and consequently subjects) exhibit longer relaxation time scales, which may be linked to brain activity approaching a critical point where the resting state could become unstable and displaying a more complex dynamics.

**Figure 6.**
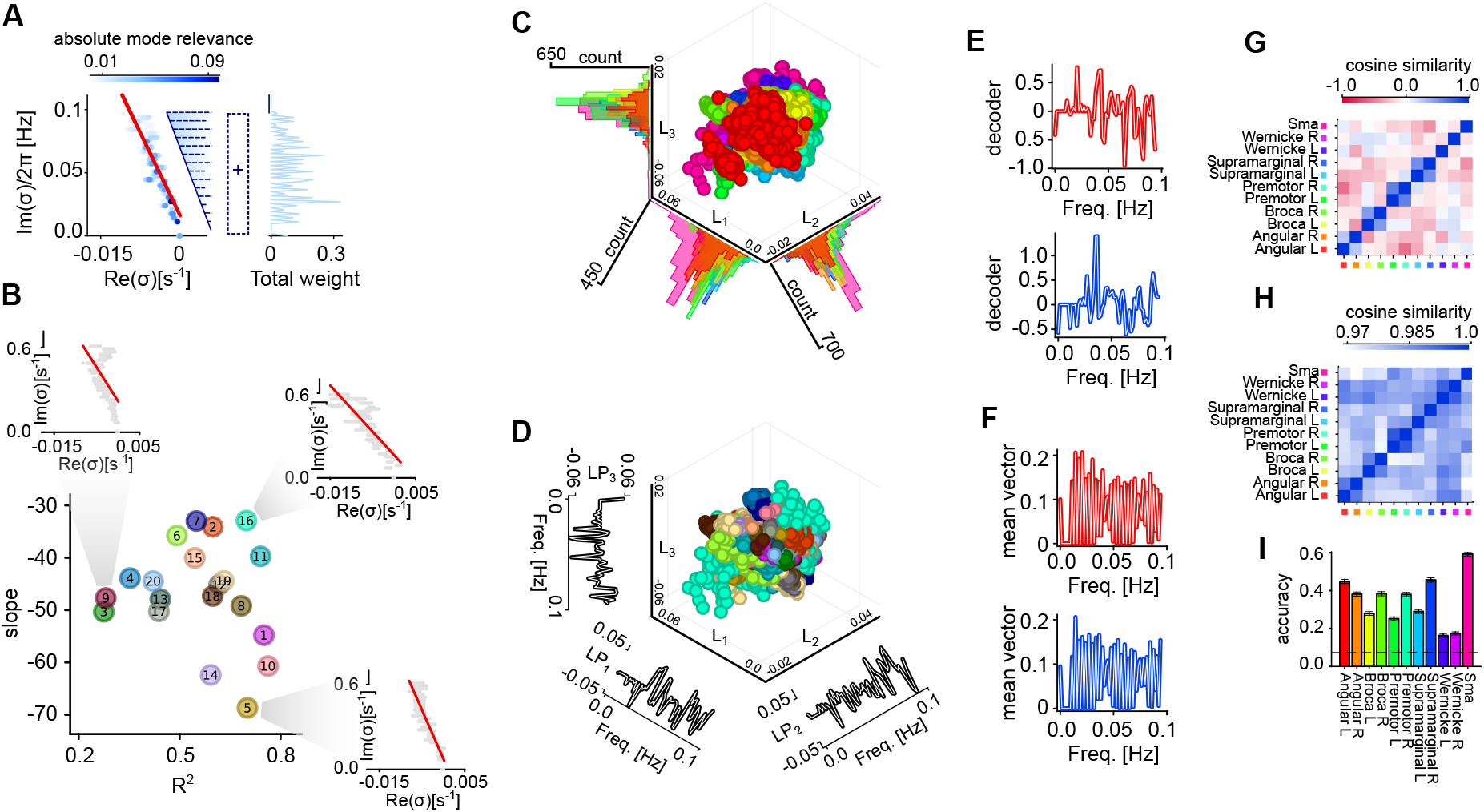
Characterization of subjects and brain areas. (**A**) Left, Spectrum of relevant eigenmodes for a sample subject. A linear regression (red line) is performed between the real and imaginary parts of the poles, *σ*_*k*_, with the slope serving as the first feature that characterizes the subject. The coefficient of determination, *R*^2^, from this regression represents the second feature. Right, Frequency vector, i.e., the histogram of Im *σ*_*k*_ each counted with multiplicity given by the averaged absolute mode relevance across voxels in a single brain area: ⟨|Ξ_*jk*_|⟩_*j∈*BA_. (**B**) Slope of linear regression and *R*^2^ of the 20 subjects in the dataset. Insets: eigenmode spectra of three representative subjects with the related linear regressions (red lines, see main text for details). (**C**) First three latent variables derived from the linear discriminant analysis (LDA) applied to the frequency vectors of 11 brain areas across all 20 subjects. The densities of circles for each latent variable are also plotted. For each subject, 100 lRNNs are inferred by sampling different endogenous noise in their stochastic dynamics, resulting in a total of 22,000 circles. Colors code brain areas according to Fig. 2A. (**D**) Same as (C) but with circles colored according with the subject color as in (B). Composition of the three latent variables LP_1_, LP_2_, LP_3_ are show on the side of their respective axes. (**E** and **F**) Linear decoders (E) discriminating the different areas in the frequency-vector space and average frequency vectors (F) for two sample brain areas (red and blue). (**G** and **H**) Cosine similarity between linear decoders (G) and between frequency vectors (H) per area averaged across subjects. (**I**) Accuracy of decoders along the training set. Dashed black line highlights the random chance level separation.

Shifting our focus to the frequency vectors derived from the inferred lRNN, we sought statistical regularities that reveal invariant features associated with individual brain areas and subjects. The frequency vector for a given brain area in a subject is represented by the histogram of Im *σ*_*k*_ from the corresponding lRNN, each counted with multiplicity proportional to its absolute mode relevance averaged across the voxels in that area: ⟨|Ξ_*jk*_|⟩ _*j∈*BA_. We then conducted a linear discriminant analysis (LDA) in the frequency-vector space to identify patterns that effectively discriminate between different brain areas (Fig. 6C). Despite originating from distinct subjects, the resulting data points clustered according to their respective brain areas. It is important to note that the compositions of the latent variables (L_1_, L_2_, and L_3_) correspond to specific frequency patterns (Fig. 6D). Among these, the pattern associated with L_1_ appears most relevant, as data points on this linear manifold are distinctly clustered based on the brain area of belonging (Fig. 6C).

To further isolate the frequencies at which each brain area exhibits unique dynamics, we computed the corresponding linear decoders (see Materials and Methods). The elements of the coefficient vectors characterizing each decoder are assigned high absolute values if the associated frequency plays a crucial role in distinguishing that area from others (Fig. 6E). The resulting patterns are markedly distinct from the average frequency vector, which, in contrast, reveals a continuum of relevant frequencies (Fig. 6F). In fact, the frequency vectors show significant similarities, making it challenging to differentiate between areas even if they are located in different hemispheres (Fig. 6H). In contrast, the decoders exhibit a lower degree of similarity, except when comparing the corresponding areas of both hemispheres (Fig. 6G). Interestingly, there is an exception to this trend: the decoder for the right Wernicke area appears to highlight frequencies that differ from those identified for the same area in the left hemisphere. Further insights into brain lateralization are revealed by examining the accuracy of the decoders. Specifically, it is more difficult to linearly recognize left hemisphere patterns by this method for the Broca, Premotor, and Supramarginal areas (Fig. 6I). It is worth noting that Wernicke is the most difficult to recognize.

In short, these findings demonstrate that inferred lRNNs can serve as powerful proxies to characterize the unique dynamics of subjects and areas of the brain, capturing their specific dynamical footprints.

## Discussion

The successful application of linear recurrent neural networks within the reservoir computing framework represents a significant methodological contribution to model resting-state fMRI data. Our results show that lRNNs effectively capture brain dynamics, suggesting that resting-state activity can be approximated as a linear system in a high-dimensional space, supporting recent studies advocating for linear models in this context (44). Our approach provides valuable and easily accessible insights into the functional connectivity structure as similarity patterns between vectors associated with the eigenmode decomposition of the inferred lRNN. This low-dimensional description of the whole brain activity moves the focus on the modes subspace offering a normative framework where brain areas and inter-subject comparison is more straightforward.

Notably, the mode dynamics described in Eq. (6) provides an equivalent linear representation for the observables of the system under investigation – namely, the single-subject voxel-wise BOLD activity. The resulting vector of all time derivatives (see Materials and Methods) serves as an alternative descriptor of the system’s state (45), potentially encapsulating the trajectory evolution at any given time. This concept is reminiscent of the dynamic mode decomposition (DMD) with Hankel matrix (46, 47), but with a significantly reduced dimensionality constraint. Unlike the DMD approach, which requires estimating a number of parameters that scales quadratically with the number *M* of observables – specifically (*M* × *T*)^2^ parameters (where *T* denotes the number of delayed time samples) – our reservoir computing approach only necessitates inferring order *N* readout weights, corresponding to the size of the lRNN. This result also stems from considering an optimal amount of memory-less fluctuations integrated into the lRNN units’ endogenous activity. This feature enables effective modeling of non-informative fast fluctuations in BOLD activity without requiring inference of additional high-frequency, fast-decaying modes, thereby reducing the dimensionality of the latent state space visited by the deterministic components of the inferred lRNNs. This characteristic makes the presented method particularly advantageous and flexible, allowing the inference of lRNN-based digital twins at the single-subject level using a limited amount of experimental data.

The reported results have the potential to open up several avenues for future research. Indeed the possibility offered to represent the single-subject as a point into a low-dimensional map – i.e., a landscape (Fig. 6B) – allows in principle to follow in time its drift in longitudinal studies like those focused on neurodegenerative diseases (48) or aging (49, 50, 51). Here, the novelty is the fully data-driven nature of the approach to build such landscape of subjects compared to those relying on model-driven inferences (52). Characterizing the trajectory followed in this landscape by each subject could provide valuable insights into the alterations in functional connectivity and dynamics. For instance, the emergence of unstable modes may be linked to transitions into pathological states of a specific neurological disease. By perturbing the system with specific and personalized rehabilitative or pharmacological treatments that aim to suppress these “pathology-related” modes, we may gain a means to intervene and reduce the occurrence of such transitions, ultimately helping to tailor optimal therapeutic strategies (28, 53).

In line with this, as in classic digital twins, the inferred lRNN can incorporate any additional available information about the system simply as new input. An external stimulation can potentially act as a selector for different dynamical regimes, allowing for two distinct representations with a single network (54). The functional connectivity itself could be a proxy for state transition. Indeed, a significant change in functional connectivity might indicate that the decomposition has also changed, suggesting that the observed system is close to a different equilibrium point (55). These changes can be tracked following the distribution of poles that characterize the autonomous dynamics of the inferred linear RNN. This is the case for transitions in global brain states – such as those governing the sleep-wake cycle – that arise from network destabilization, a hallmark of criticality (56, 57, 58). Under these conditions, pole distributions correlate with longer relaxation time scales and steeper slopes (as illustrated in Fig. 6B), possibly underlying previously observed state-dependent spectral signatures (59). This metastable dynamics results in broader excursions within the latent state space of neural activity, making predictions of future BOLD activity more challenging for the inferred lRNN. The quality of lRNN predictions can then serve as a proxy for dynamical stability, akin to findings in intracortical local field potentials (LFPs) of macaque monkeys during anesthesia-induced transitions between wakefulness and unconsciousness (60).

While this study demonstrates the efficacy of lRNNs in modeling resting-state brain activity, several limitations must be acknowledged. The assumption of linearity, although supported by our results, may not fully capture the complexity of brain dynamics under all conditions. Nonlinear phenomena, transient states, and the influence of external stimuli could require more sophisticated modeling approaches (61, 14). Alternatively, the apparent suitability of resting-state activity being well represented by a stationary multivariate Ornstein-Uhlenbeck process (i.e., Brownian motion) (62, 63) may reflect an intrinsic limitation of the BOLD signal in conveying detailed neural dynamics. Indeed, this hemodynamic-related signal acts as a lumped observable that inevitably linearizes the spiking activity of relatively large neuronal assemblies (64, 44). The coexistence of endogenous noise with the relatively high-dimensionality of the BOLD time series can give rise to a multivariate Brownian motion exhibiting a rich repertoire of restless spatiotemporal modes (63). These are captured in our lRNN by relatively wide distributions of poles. Within this linear stochastic system, noise spontaneously excites its normal modes, eliciting co-fluctuating patterns closely linked to the functional connectivity measured in rs-fMRI. Interestingly, this structured stochasticity may, at least in principle, exhibit similarities to turbulence (65, 66).

However, the existence of a single fixed-point around which all voxel activities fluctuate, imposes a rather strong constraint on the brain’s ability to perform information processing. Indeed, this computational capability is thought to arise from the complex unfolding of neural trajectories across a landscape rich in saddles and metastable states (67, 68). Yet, lRNNs cannot represent the latent state space of a multistable system. This raises an important question: how can we reconcile the evidence that resting-state fMRI activity is well described by stochastic lRNNs with the necessity of nonlinear dynamics for brain computation? A tentative answer is that, during rest, endogenous fluctuations have a magnitude comparable to the deterministic excursions of neural activity associated with relaxation dynamics. Consequently, the nonlinear components of brain activity may only become apparent when the system is pushed far from equilibrium – such as when the brain engages in cognitive functions like motor or perceptual decision-making. A promising direction for future work would then be to test this hypothesis by investigating to what extent an effective lRNN can be inferred from BOLD time series recorded while subjects engage in such cognitive tasks.

## Materials and Methods

### Dataset description

#### Subjects

The study included 20 healthy, right-handed subjects (mean age ± SD = 37 ± 12; 7 females, 13 males) with no history of neurological disorders. The study was approved by the Institutional Review Board at Memorial Sloan Kettering Cancer Center, and informed consent was obtained from each participant. During the resting-state condition, subjects were instructed to lie in the scanner, keep their eyes open, try to think of nothing in particular, and maintain fixation on a central cross on the screen.

#### MRI methods

A GE 3T scanner (General Electric, Milwaukee, Wisconsin, USA) and a standard quadrature head coil was employed to acquire the MR images. Functional images covering the whole brain were acquired using a (T2*)-weighted imaging sequence sensitive to blood oxygen level-dependent (BOLD) signal (repetition time, TR/TE = 2500/40 ms; slice thickness = 4.5 mm; matrix = 128 × 128; FOV = 240 mm; volumes = 160). Functional matching axial T1-weighted images (TR/TE = 600/8 ms; slice thickness = 1 mm) were acquired for anatomical co-registration purposes.

#### Data preprocessing

Functional MRI data were processed and analyzed using the software program Analysis of Functional NeuroImages (AFNI; Cox, 1996). Head motion correction was performed using 3D rigid-body registration. The first volume was selected to register all other volumes. The first volume was chosen because it was acquired before the anatomical scan. Both task and resting state fMRI scans were monitored using a real time post-processing software BrainWave (BrainWave RT, Medical Numerics, Germantown, MD) to monitor brain activity and the head motion. For subjects showing severe head motion over time, generally, the scan is repeated. For small head motion (less than 2 voxel size), a motion correction algorism (iterated linearized weighted least squares) considering 3 translation and 3 rotation parameters against a reference volume was applied. The obtained 6 parameter time courses were also integrated in the statistical analysis to regress out residual motion-correlated artifactual voxels. Spatial smoothing was applied to improve the signal-to-noise ratio using a Gaussian filter with a 6 mm full width of half maximum. Corrections for linear trend and high-frequency noise were also applied. Resting-state data requested some more preprocessing steps. They were corrected for head motion by regressing head motion data and the first five principal components of the white matter and CSF signals. They were also detrended, demeaned, and band-pass filtered (frequency range 0.01-0.1 Hz). All fMRI data were registered to the standard space (Montreal Neurological Institute MNI152 standard map).

### fMRI dimensionality analysis

Principal Component Analysis (PCA) is a widely used dimensionality reduction technique used to transform a dataset into a lower-dimensional space while preserving most of the information in the original data (69). In the context of the given problem, PCA is applied to the set of BOLD signals to reduce redundancy and keep only the relevant information.

The PCA-based dimensional reduction of BOLD is carried out based on the initial 280 s (learning period) of each subject’s data. Referring to Fig. 2, the number of principal components taken into account are those capturing 99.5% of the variance in the time series of each area. The resulting transformation is represented by a block matrix ***P***_0_, where each block’s columns ***p***_*i*_ correspond to the eigenvectors of the covariance matrix of the respective cortical area:

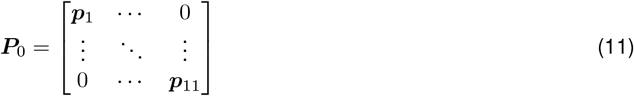

The output undergoes orthogonalization through an additional PCA step (***P***_1_) utilizing all components. Subsequently, normalization is performed by applying the diagonal matrix ***N***, which contains the reciprocals of the maximum values of the compressed signals observed during the learning period. This approach ensures that all currents enter the reservoir with the same maximum strength. The final transformation matrix ***P*** is computed as:

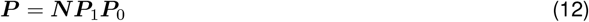

While the antitransformation matrix ***P*** ^*−*1^ is calculated as:

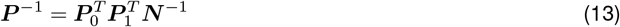

The data is effectively reduced and then restored, ensuring the preservation of essential information while reducing dimensionality for the analysis. The decomposition for voxel activity is derived by applying ***P*** ^*−*1^ to the decomposition matrix for the actual network inputs.

### Koopman Operator

The discrete-time Koopman operator 𝒦^1^ : ℱ → ℱ is a linear operator defined in an infinite-dimensional space ℱ of observables of the system state (70, 19, 20). The operation of the Koopman operator on an observable *g* is described by the equation:

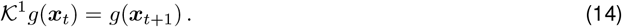

Here, ***x*** represents a point in the state space. The eigenfunctions of the Koopman operator are potential observables of the system themselves, and have the peculiar property of evolving linearly in time:

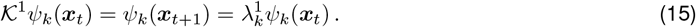

These eigenfunctions *ψ*_*k*_(***x***_*t*_) constitute a basis of the space ℱ, enabling the representation of all the possible observables of the system as a linear combination of linearly evolving functions.

In general there exists a family of Koopman operators, 𝒦^*t*^, that advances a function forward by a time *t*:

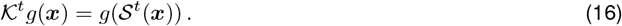

Here, 𝒮^*t*^ denotes the time flow operator, with ***x***(*t*) = S^*t*^[***x***(0)]. The infinitesimal generator of the Koopman operator family 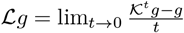 is the Lie operator which evaluates the temporal change of observables:

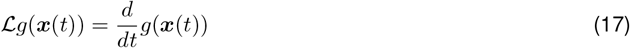

The family of Koopman operators can then be expressed in this term 𝒦^*t*^ = *e*^*ℒt*^.

### Reservoir computing

The reservoir computing approach involves a RNN with fixed and random internal couplings (namely the ‘reservoir’) whose state is fed forward to a second set of ‘readout’ units (31, 4). In this framework the required ‘echo-state property’ spontaneously emerges from the RNN collective dynamics. The units composing the RNN are intended to model homogeneous and local neuronal assemblies of a cortical network (71, 72, 73). Each of the *N* units in the network has activity state *r*_*j*_(*t*) evolving in time as

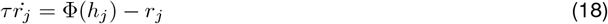

with *j* ∈ [1, *N*] ⊂ ℤ. The decay time constant *τ* and the sigmoidal activation function Φ(*h*_*j*_) is the same for all units. Here we consider the limiting case of weak recurrent coupling where the activation function can be approximated to a linear function: Φ(*h*_*j*_) = *h*_*j*_. The synaptic input *h*_*j*_ is the weighted sum

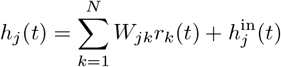

where *W*_*jk*_ are the elements of the synaptic matrix (i.e., the internal couplings) ***W* ∈** ℝ^*N ×N*^, and 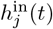 is the external input received by the unit *j*. In reservoir computing, the external input is driven by be the measured observable *ω* of the inspected system, which is fed into the network using random synaptic couplings: 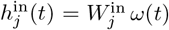. Due to the high dimensionality of the trajectories in the RNN state space, a simple linear transformation represented by ***W*** ^out^ is often sufficient to accurately map the RNN state into a desired output time series. More precisely, given ***R*** ∈ ℝ^*N ×T*^ whose *k*-th row represent the time series of the *k*-th unit during a learning period lasting *T* time steps, and **Ω** ∈ ℝ^*M ×T*^ has rows given by the *M* observables *ω*_*k*_(*t*) to be replicated, the linear map is computed by a ridge regression:

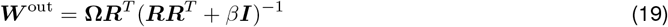

where *β* is a regularization parameter and ***I*** ∈ ℝ^*N ×N*^ the identity matrix. The target output coincides with the external input. Then, the resulting linear map can be used to simulate the external stimulation and predict the future steps of the data. This is equivalent to update the synaptic matrix of the reservoir. In particular ***W*** ^in^***ω***(*t*) ≈ ***W*** ^in^***W*** ^out^***r***(*t*) leading to the autonomous system:

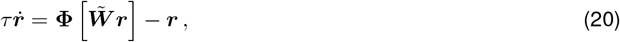

where 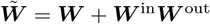 is the updated synaptic matrix.

### Linear recurrent neural networks (lRNN) and autoregressive models

In the established framework of reservoir computing with linear RNNs (lRNN, i.e., with a linear activation function) by setting as initial condition *r*_*k*_(−∞) = 0, the system Eq. (18) has the following solution:

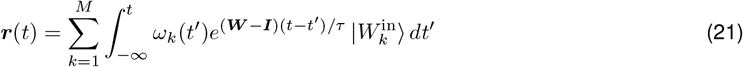

As above *ω*_*k*_(*t*) denotes the *k*-observable measured from the inspected system. Assuming a time *t*^*∗*^ exists after which a matrix ***W*** ^out^ maps the network state to the input data ***ω***(*t*) = ***W*** ^out^***r***(*t*), we can rewrite the above equation as:

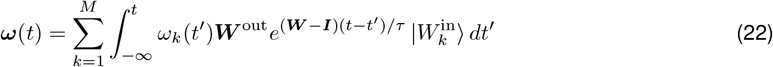

By substituting *s* = *t* − *t*^*′*^, we obtain an autoregressive description of the signals:

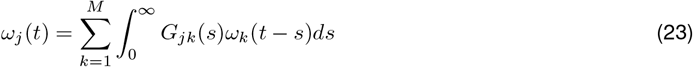

Here, 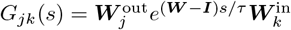. These kernels can be expressed in terms of the synaptic matrix spectrum. The diagonal representation of the exponential matrix is given by:

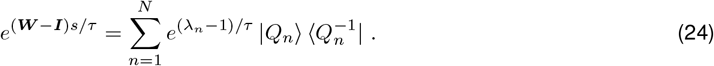

In this representation, *λ*_*n*_ denotes the eigenvalues and 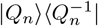 the outer product between the right and left eigenvectors of ***W***. This representation allows us to rewrite the kernel as the weighted sum over exponentially

decaying components:

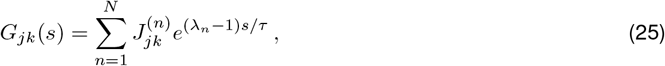

where the tensor of weights is defined by the set of rank-1 matrices 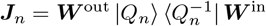.

### lRNN and the Laplace transform of system state

Assuming the existence of a time *t*^*∗*^ such that a matrix ***W*** ^out^ maps the lRNN state into the system observable given as input *ω*(*t*) = ***W*** ^out^***r***(*t*), the open-and closed-loop dynamics are in principle equivalent (31, 30). This property allows us to link the kernel of the autoregressive model (closed-loop dynamics) to the spectrum of the learned synaptic matrix 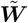, and consequently, to the resulting finite Koopman approximation, as we will see in the following.

#### Open-loop Representation

The autoregressive formula expressed in Eq. (3) can be reformulated as:

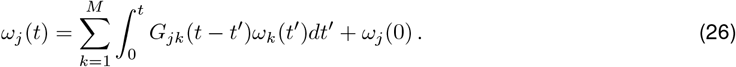

The convolution theorem for the Laplace transform leads to:

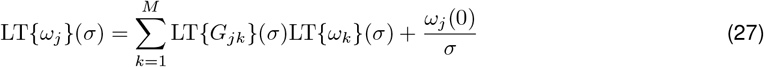

Here, 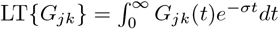 denotes the Laplace transform of the kernel that is

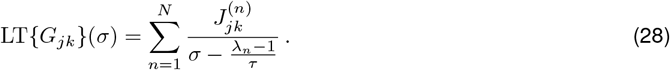

Thus, in matrix formalism the Laplace transforms of the system observables result to be

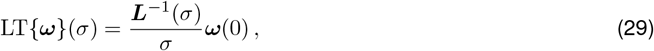

where the matrix ***L***(*σ*) ∈ ℝ^*M ×M*^ has elements *L*_*ij*_(*σ*) = *δ*_*ij*_ − LT{*G*_*ij*_}(*σ*).

#### Closed-loop representation

The closed-loop network evolves according to Eq. (5). Assuming a diagonalizable synaptic matrix 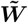, it can be factorized as 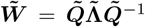, where 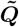 is a matrix whose columns are the right eigenvectors of 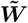 and 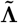 is a diagonal matrix containing the associated eigenvalues. The spectral decomposition of the learned synaptic matrix leads to a decoupled set of linearly evolving projections 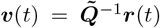 whose *N* elements evolves independently as

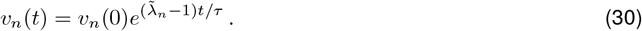

These variables serve as a basis to decompose the system observables *ω*_*k*_(*t*) as the sum of linearly evolving modes, resembling the Koopman decomposition. Indeed, given 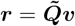 we can write

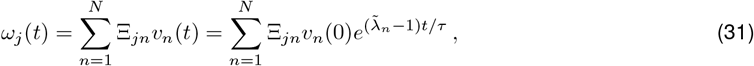

with matrix **Ξ** ∈ ℝ^*M ×N*^ defined as **Ξ** = ***W*** ^out^***Q*** such that ***ω***(*t*) = **Ξ*v***(*t*). The Laplace transform of the replicated observables is then

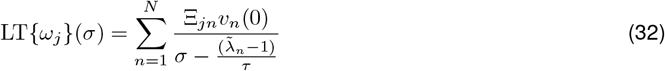

whose poles coincide with the eigenvalues of the estimated Koopman matrix.

Given the equivalence between the open and closed representations, *L*_*ij*_(*σ*)*/σ* shares the same poles, providing a link between the autoregressive modeling and the linear dynamic representation.

### lRNNs and the spectrum of the Koopman operator

The closed-loop representation provides a decomposition of the signal as linear evolving modes. We can explicitly express the functional form of the basis starting from open loop dynamics. By applying 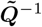 to both hand sides of Eq. (2) yields:

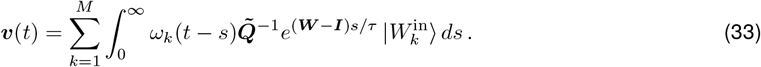

By resorting to the spectral decomposition in Eq. (24), the above eigenmode dynamics reduces to:

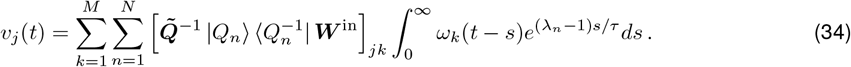

We can then expand in Taylor series the observables: 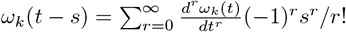. This allows to solve the above integral leading to a functional description of the estimated eigenmodes

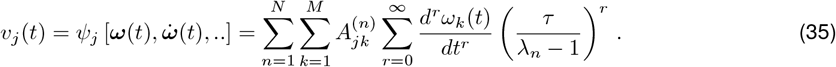

The vector of all the derivatives *d*^*r*^*ω*_*k*_(*t*)*/dt*^*r*^ fully determine the state of the observed system (33) and *ψ*_*j*_ is the eigenfunction of the approximated Koopman operator depending on the full state of the system. The corresponding eigenvalue 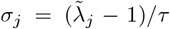, and 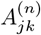 is a tensor of complex weights defined by the rank-1 matrices 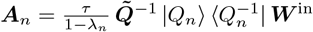.

### Stochastic lRNN

The deterministic neural network described can be extended to the stochastic case, where a white noise input stimulates the reservoir, maintaining it out of equilibrium as a continuous source of new energy. Each unit receives a total input defined by the equation:

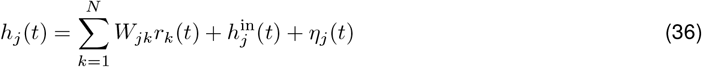

where *η*_*j*_(*t*) represents an Ornstein-Uhlenbeck process (colored noise) with a relatively small correlation time such that for the purpose of this work it can be considered as memory-less: ⟨*η*_*i*_(*t*)*η*_*j*_(*t*^*′*^)⟩ = *γ*^2^*δ*_*ij*_*δ*(*t ™ t*^*′*^). An optimal standard deviation *γ* can be identified to maximize the accuracy of the linear map ***W*** ^out^.

In the closed-loop formulation, the input can be expressed as:

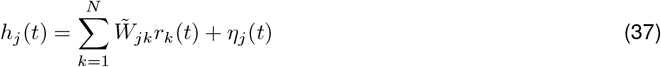

Consequently, the associated Langevin equation for the reservoir is given by:

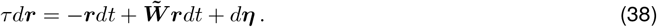

In this context, the introduced autoregressive description and linear decomposition remain valid when considering the expected values of the system.

### Functional Connectivity and Functional Connectivity Dynamics

Functional Connectivity (FC) is a measure of the degree of co-activation over time of different brain regions. Although FC does not allow to infer the directional flow of information among the nodes of the brain network, it has proven to provide valuable insights into the functional organization of the central nervous system (74). Mathematically, the FC between two brain regions labeled A and B, is usually measured as the correlation function

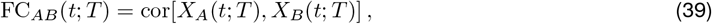

where cor is the Pearson correlation coefficient, and *X*_*A*_(*t*; *T*) and *X*_*B*_(*t*; *T*) are the BOLD signals from region A and B, respectively, in the time window [*t, t* + *T*].

Temporal changes of functional connectivity are evaluated by computing the Functional Connectivity Dynamics (FCD) (75, 76). FCD is defined as the correlation between vectorized FC matrices evaluated in time-shifted chunks of BOLD activity:

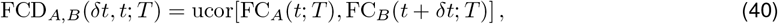

where *δt* is a constant time shift, and ucor is the correlation between the vectors composed of the upper-diagonal elements of the two FC matrices involved.

Throughout the whole paper FC is computed with *t* = 0 s and *T* = 387.5 s, while FCD uses *T* = 75 s time shift *δt* multiples of 12.5 seconds.

### Functional Connectivity in lRNN

Once a lRNN capable to reproduce the systems observables *ω*_*j*_(*t*) is available, the elements of the FC matrix can be inferred directly from the lRNN parameters. The computed Functional Connectivity of the simulated signals depends on the covariance matrix estimated from time *t* over a time window of length *T*. Considering large time windows *T* and assuming a zero time average ⟨*ω*_*j*_(*t*)⟩_*t*_ ≃ 0, it holds:

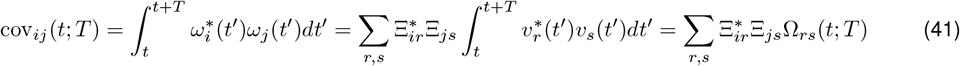

Here we are making use of the spectral decomposition in Eq. (7) and we define Ω_*rs*_(*t*; *T*) as the estimated covariance between the OU modes *v*_*r*_, *v*_*s*_. From the simulations, it results that only a few slow uncorrelated modes have a significant weight Ξ in the decomposition. In this scenario, the covariance matrix is well approximated by the dot product between the representational vectors in the functional space:

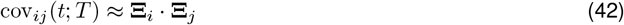

Consequently for the correlation matrix it holds

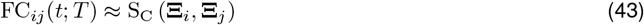

where S_C_ indicates the cosine similarity function, i.e., the Pearson correlation between row and column vectors composing **Ξ**^*∗*^ and **Ξ**, respectively, under the assumption of a zero mean for the system observables.

#### Simulation parameters

Network parameters have been selected through grid search methods to optimize prediction accuracy. The reservoir state is initialized in a zero-activity state and is disrupted from equilibrium by the input for a transient period of 20 *s* before learning. This period corresponds to the *t*^*∗*^ mentioned in the preceding paragraphs. The input signal is linearly interpolated, and the system’s dynamics are resolved using the Euler–Maruyama method with a time step that is one-hundredth of the sampling interval. The spectral radius *ρ* of the synaptic matrix, used for fMRI analysis, is set to 0.5, and the time constant *τ* is 1.0. The standard deviation *g*^in^ of the input matrix ***W*** ^in^ is set to 1. The thermal noise shown in Fig. 3 has a regularizing effect. Consequently, we do not employ further regularization for the readout regression. For the example system depicted in Fig. 1, the parameters are *ρ* = 0.5, *τ* = 0.1 with no regularizers, and *g*^in^ equal to the inverse of the square root of the number of inputs. The stochastic scenario is characterized by thermal noise with a standard deviation of 10^*−*5^ and ridge regression with a regularization of 10^*−*7^. For cases with missing variables, the parameters are set to *ρ* = 0.6, *τ* = 0.25.

### Clustering analysis

The clustering analysis is performed on the resulting spectrum of the data by discretizing the complex space into a grid with a real resolution of 0.00025*s*^*−*1^, and an imaginary resolution of 0.01*s*^*−*1^.

To further reduce the dimensionality of the data and get an interpretable 3 dimensional representation, Linear

Discriminant Analysis (LDA) is employed. LDA is a widely used technique in machine learning (69). It is a supervised learning method that aims to reduce the dimensionality of the data while preserving the discriminatory information between different classes. The core idea is to find a linear transformation that maximizes the ratio of the between-class scatter ***S***_*b*_ to the within-class scatter ***S***_*w*_. This optimization process equals to solve the following generalized eigenvalue problem

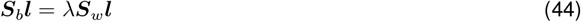

The optimal projection matrix ***L*** to a subspace of dimension *k* is given by the eigenvectors ***l*** corresponding to the largest *k* eigenvalues.

### Linear Decoders

To find the distinct frequency patterns that characterize each brain area, we computed the associated linear decoder within the discretized frequency-vector representation. This process involved linking each brain area to a target versor in an 11-dimensional space. Then we employed a pseudoinverse algorithm to derive the decoders, which maps the frequency-vectors – resulting from all subjects and 100 reservoir realizations – to their corresponding relative versors.

### Manuscript Language Optimization

To enhance the clarity, coherence and readability of the manuscript, we employed large language models (LLMs) for language editing and refinement. All edits assisted by the model were subsequently reviewed by us to ensure accuracy and to preserve the intended scientific meaning.

## Acknowledgments

We thank M. Allegra for the feedback on an earlier version of the manuscript and S. P. Caminiti, I. Branchi and G. V. Vinci for the stimulating discussions. Work partially funded by the NEXTGENERATIONEU and MUR (PNRR- M4C2I1.3) project MNESYS (PE0000006–DD 1553 11.10.2022) and project EBRAINS-Italy (IR0000011– DD 101 16.6.2022) to MM.

